# A comparison of high resolution image reconstruction methods from confocal array detector with a new deconvolution algorithm

**DOI:** 10.1101/2021.08.02.454749

**Authors:** Sylvain Prigent, Stéphanie Dutertre, Aurélien Bidaud-Meynard, Giulia Bertolin, Grégoire Michaux, Charles Kervrann

## Abstract

Array detector allows a resolution gain for confocal microscopy by combining images sensed by a set of photomultipliers tubes (or sub-detectors). Several methods have been proposed to reconstruct a high resolution image by linearly combining sub-detector images. To overcome the limitations of these techniques, we propose a new reconstruction method that takes the full stack of spatially reassigned detector signals as input. We show on both calibration slides and real data that our deconvolution method allows to achieve a better reconstruction performance in terms of resolution, image contrast, and spatial intensity homogeneity. The tested algorithms are available in an open source software.

---

The principle of array detector confocal microscopy is to replace the point detector by a set of sub-detectors circularly arranged around a central detector [1] (Fig. 1). Accordingly, the optical resolution is no longer determined by the size of a single pinhole but depends on the relative positioning of sub-detectors, each acting as a small pinhole and capturing a slightly different view on the sample. With this array detector approach, more light is collected and the reconstructed image has a higher signal-to-noise ratio (SNR). Nevertheless the estimation of the high resolution image from sub-detectors is not straightforward. Since the emergence of this technology, two categories of methods have been investigated for image reconstruction: i) the sub-detector combination methods [2–6] allow either a gain of contrast or a gain of resolution but not both; ii) the deconvolution-based methods are more sophisticated and allow both contrast and resolution gain, but may create spurious artifacts.

**Fig. 1.**
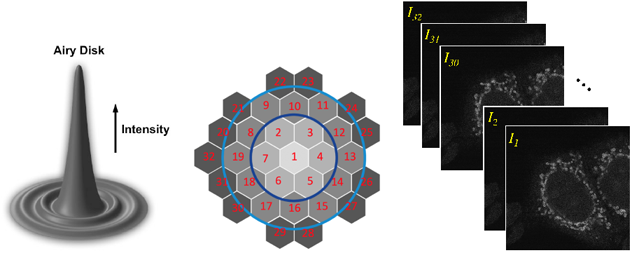
Scheme of the array detector and image stack. The 32 sub-detectors are hexagonal and arranged around the central sub-detector. The dark blue (resp. light blue) circle highlights the group of *N* = 7 (resp. *N* = 19) central detectors. The detectors inside/outside the circles are called “*inner*”/”*outer*” detectors.

In this Letter, we first review the sub-detectors combination techniques and we describe a data-driven approach for automatically estimating the weighting parameters. Second, we propose an original deconvolution algorithm that estimates a single deconvolved image directly from the array of low/high frequency images. We show that it allows us to achieve a better compromise in terms of resolution and signal reconstruction when compared to existing methods.

The array detector we consider consists of 32 sub-detectors arranged around a central sub-detector as illustrated in Fig. 1. Each sub-detector has a diameter of 0.2 AU (Airy Unit). The first inner ring grouping the *N* = 7 central detectors has a diameter of 0.6 AU, the second inner ring grouping the *N* = 19 detectors has a diameter of 1 AU, and the full detector has a diameter of 1.25 AU. Among the list of techniques that were analyzed, the base-line method (pseudo-confocal (PC)) amounts to summing *N* images as follows:

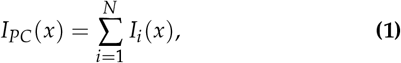

where *I_i_*(*x*), illustrated in Fig. 1 (right), denotes the fluorescence intensity observed at the spatial position *x* ∈ Ω (Ω is the image domain) sensed by the sub-detector with index *i*. If *N* = 7, 19 and 32, the resulting images are equivalent to confocal images with a pinhole at 0.6 AU, 1 AU, and 1.25 AU, respectively. Note that summing the *N* images enables to achieve an image with a higher contrast than a conventional confocal image. Since, image scanning microscopy (ISM) technique [2, 3] has been proposed to produce a higher resolution image. The principled approach consists in reassigning the signal from each sub-detector to the central detector as follows:

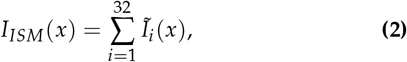

where *Ĩ_i_*(*x*) is the intensity at pixel *x* sensed by the sub-detector *i*, spatially co-registered to the central detector *i* = 1. Later on, virtual fluorescence mission difference (FED) [4, 5] has been defined to create an high resolution image. Conventional FED microscopy scans twice the sample, first with a point shape illumination, and second with a donut shape illumination. Virtual FED is similarly obtained by subtracting the “outer” ring of sub-detectors to the “inner” sub-detectors:

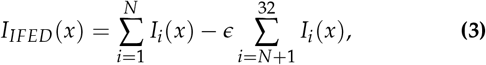

where *∊* > 0 is a subtraction factor and *N* is set to 7 or 19, depending on the compromise made between intensity contrast and spatial resolution. Finally, Li *et al*. [6] showed that, by substituting the ISM image (2) to the sum of the *N* row detectors as follows:

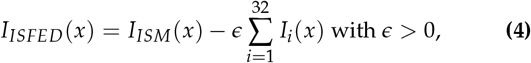

we can obtain a gain of about 12% in terms of resolution can be achieved when compared to the original IFED method.

The performance of the so-called ISFED and IFED methods depends on the calibration of the *e* parameter in (3)–(4). In [5], the authors showed that by setting *∊* = 0.3 yielded very satisfying experimental results. Nevertheless, we experimentally observed that *e* should be adaptively adjusted according to SNR; for low SNR values *∊* = 0.3 is appropriate, but *∊* must be in-creased to 1.0 to accommodate high SNR values. These results suggested the development of the following data-driven method [7] to automatically set *∊*. As starting point, we considered the following general formulation^1^ based on (3) and (4):

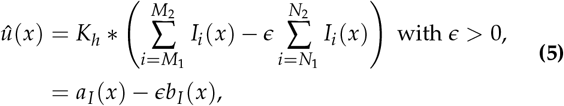

where *K* refers to a 2D spatial convolution filter with a kernel size *h*, and * denotes the convolution operator. In (5), 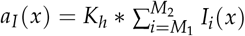 is the sum of the detectors with indexes between *M*_1_ and *M*_2_, and 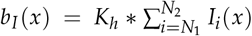 is the sum of the detectors with indexes between *N*_1_ and *N*_2_. The kernel size *h* is chosen very small to both slightly reduce noise and preserve high frequencies. In [7], we considered the following local Stein’s unbiased risk estimate (SURE) to optimally determine *∊*:

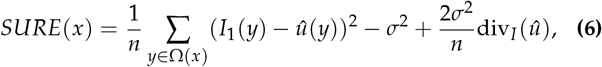

where Ω(*x*) is a local spatial neighborhood centered at pixel *x*, *n* is the number of pixels in Ω(*x*) (i.e., *n* = |Ω(*x*)|) assumed to be constant for all pixels in Ω, *σ*^2^ denotes the variance of the assumed Gaussian noise, and div is the divergence of *û* wrt 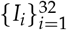. Furthermore, we defined the empirical SURE as the sum of local SURE: *SURE* = ∑_x∈Ω_ *SURE*(*x*). Solving 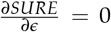 gives the closed-form solution (see [7], Supplementary Note):

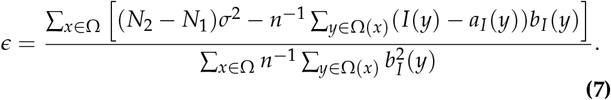

The ISM, IFED and ISFED techniques are appealing as they allow to improve resolution with simple calculus, without losing too much frequency information. Nevertheless, these linear methods are limited as the input images can contain subtle structural details and low weak signals. To overcome the current limits of ISM, IFED, and IFSED methods, conventional deconvolution algorithms such as Richardson-Lucy or Wiener that take the ISM image as input, have been investigated in array detector microscopy. In the remainder of the Letter, we will denote ISM-RL (Richardson-Lucy) and ISM-W (Wiener) the two corresponding methods. However, deconvolving the ISM image is sub-optimal as the input image results from the merging of *N* sub-detectors inducing the loss of subtle structural information and high frequencies. As a remedy to this problem, we propose to apply a performant deconvolution algorithm (SPITFIR(e) [8]), directly on the native array data (that is the array detector). In summary, SPITFIR(e) amounts to finding the image *u* that best minimizes a global energy *E*(*u*, *f*) composed of a data fidelity term and a sparse-promoting regularization term *R_ρ_*(*u*):

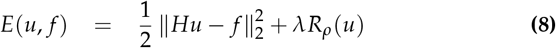

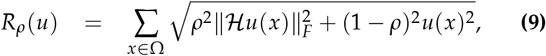

where *f* is the noisy and blurred image, *H* represents the point spread function (PSF) (matrix form), *u* is the restored image, and *λ* is a regularization parameter. In (9), *ρ* ∈ [0, 1] involved in the sparse-promoting regularizer, is a weighting parameter that balances the Hessian term 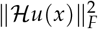 used to encourage smooth variations of the signal and the intensity term *u* that “weakly”, “moderately” or “strongly” encourages sparsity in the restored image (see details in [8]). In practice, SPITFIR(e) was successfully used for restoring multi-dimensional fluorescence microscopy images corrupted by Poisson-Gaussian noise and acquired with spinning disk confocal, STED, and multifocus microscopy. Here, SPITFIR(e) is applied to the ISM image (i.e., *f* = *I_ISM_*) to generate a deconvolved image (denoted ISM-S (SPITFIR(e))). Nevertheless, this energy can be modified as follows to take the 32 sub-detectors as input:

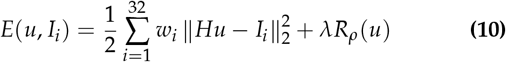

where *w_i_* = *Z*^−1^ exp(−*d_i_*/*τ*) > 0 is a weight and *Z* is a normalization constant such that 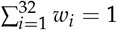. We define *d_i_* as follows: *d_i_* = *d_kl_* if *i* ∈ {*k*,…, *l*} where *d_kl_* is the averaged Euclidean distance (in pixels) of sub-detectors with indexes {*k*,…, *l*} to the central detector. We considered four groups of sub-detectors as illustrated in Fig. 1: {1}, {2,…, 7}, {8,…, 19}, and {20,…, 32}. By setting *τ* = 3.0, twice more importance is given to (high-frequency) “outer” sub-detectors {20,…, 32} than to “inner” detectors {2,…, 7}. Thus, the so-called AD-S (Array Detector - SPITFIR(e)) algorithm performs both the merging of multiple images {*I*_1_,…, *I*_32_} and deconvolution at once.

In our experiments, we used the SURE criterion to select an optimal value *∊* (see (7)) to generate the ISFED and IFED images. The underlying parameters (e.g., *λ*) of SPITFIR(e) are automatically estimated from the noisy images as described in [8]. The size of the PSF in (10) is selected in the range [1.0,…, 2.0] pixels and set to 1.5 pixels by default. We applied the eight reconstructions methods PC, ISM, IFED, ISFED, ISM-RL, ISM-W, ISM-S and AD-S to an image depicting ring shape object (“donut”) taken from the Argolight calibration slide, whose surface is known to be spatially smooth. Figure 2(a) shows the high resolution images obtained with each method. First, we can notice that PC and ISM have high SNR and low resolution. The IFED and ISFED images have low SNR due to sub-detector subtraction, but a higher resolution than PC and ISM. At first glance, the three deconvolution methods ISM-RL, ISM-W and ISM-S produced higher resolution images. Nevertheless, they are sensitive to small intensity variations, and consequently the intensity of the donut is not homogeneous. Finally, the AD-S algorithm produced an image where the ring that appears to be visually thinner and more homogeneous than the images resulting from the deconvolution of the input ISM image. In each case, we plotted 12 profiles across the donut separated by an angle of 15 degrees each (Fig. 2(b)). We can notice that the peaks corresponding to PC and ISM are large and noisy. From the conventional full width at half maximum (FWHM) criteria, the average thickness of the donut is 323 nm and 313 nm with the PC and ISM methods, respectively. The profiles obtained with IFED and ISFED methods appear to be very noisy. Nevertheless, the peaks are thinner than before, and the average thickness of the donut is 278 nm and 280 nm for IFED and ISFED, respectively. The deconvolution methods that take the ISM image as input produced high resolution images with thinner and smoother peaks. The resulting average thickness are 257 nm (ISM-RL), 275 nm (ISM-W), and 254 nm (ISM-S). Finally, the AD-S algorithm provided the best results; the profiles are less noisy, thinner (average thickness: 252 nm) and show a slightly lower variance around the donut that the other deconvolution methods.

**Fig. 2.**
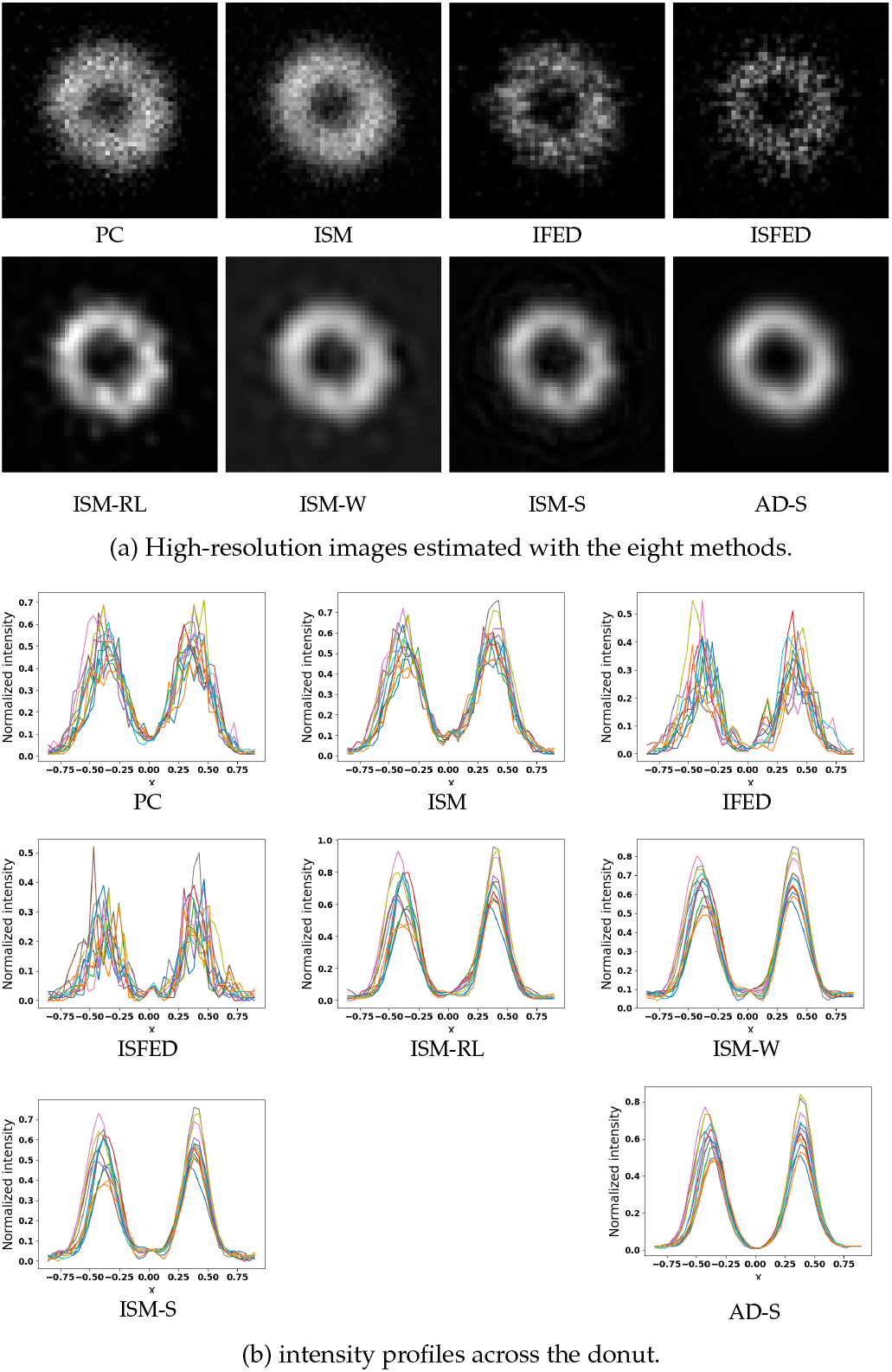
High-resolution reconstruction obtained with the eight methods applied to the “donut” image from the Argolight calibration slide (ZEISS Airyscan microscope). (a) high resolution images. (b) Twelve profiles are plot across the donut taken at 15 degrees from each other. The *x* axis represents the distance in microns to the ring center.

Furthermore, we applied the eight reconstruction methods to an image depicting vertical line pattern from the Argolight calibration slide. This pattern contains ten groups of lines where the distance between lines decrease from 550 nm to 100 nm with 50 nm steps (see Figure 3(a)). To estimate the spatial resolution, we measured the intensity gap (contrast) between the maximum and minimum intensity profile for each line. With this Argolight sample, it turns out that a contrast below 26.5% means that objects at the line resolution cannot be distinguished. The measured contrasts are plotted in Fig. 3(b). We got two groups of methods, the methods including PC, ISM, and IFED that cannot distinguish the lines closest to 350 nm and the methods that can distinguish the lines until 250 nm. Note that the ISFED image has a resolution as high as those obtained with the deconvolution methods, but is more noisy.

**Fig. 3.**
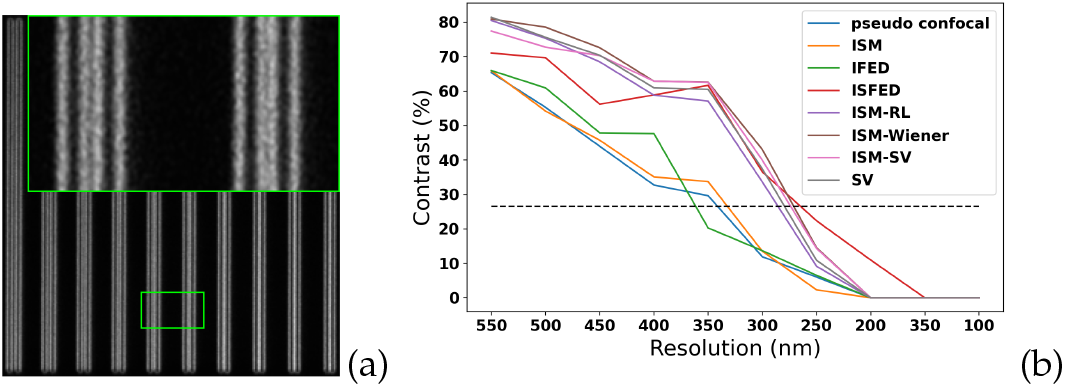
Resolution measurement using the vertical line pattern. (a) image of the pattern. From left to right, lines are spaced by 550, 500, 450, 400, 350, 300, 250, 200, 150, and 100 nm. (b) Contrast in percentage of intensity between maximum intensity and minimum intensity for each of the 10 lines patterns profiles. The black dashed line shows the resolution measurement threshold.

To evaluate the performance on real data, we first applied the eight reconstruction algorithms to an image depicting mitochondria in MCF7 cells expressing mito-GFP. In Fig. 4(a), ISFED and the deconvolution methods provided the best results in terms of resolution. However, the SNR is lower with the ISFED method. Moreover, AD-S is more robust to noise than the other competing deconvolution methods, and the resulting image has a higher resolution. If we compare the intensity profiles along the yellow lines drawn in Fig. 4(a) and estimated by each method (see Fig. 4(c)), we can notice that ISFED that produced the thinner profiles but this profile is more noisy than the others. As before, AD-S gives the best compromise between SNR and resolution, followed by ISM-S, ISM-RL, ISM-W and further IFED, ISM and PC. Furthermore, we applied the reconstruction methods to an image depicting intestinal microvilli from an adult *C. elegans* worm expressing the ERM-1::mNeonGreen fusion protein (ERM-1 is specifically localized in intestinal microvilli, see Fig. 4(b)). The ISFED and deconvolution methods better enhance the 100 nm large microvilli. Moreover, in Fig. 4(d), the cross-section profiles along the yellow lines drawn in Fig. 4(b) suggest that AD-S better removes the background (deeper curve).

**Fig. 4.**
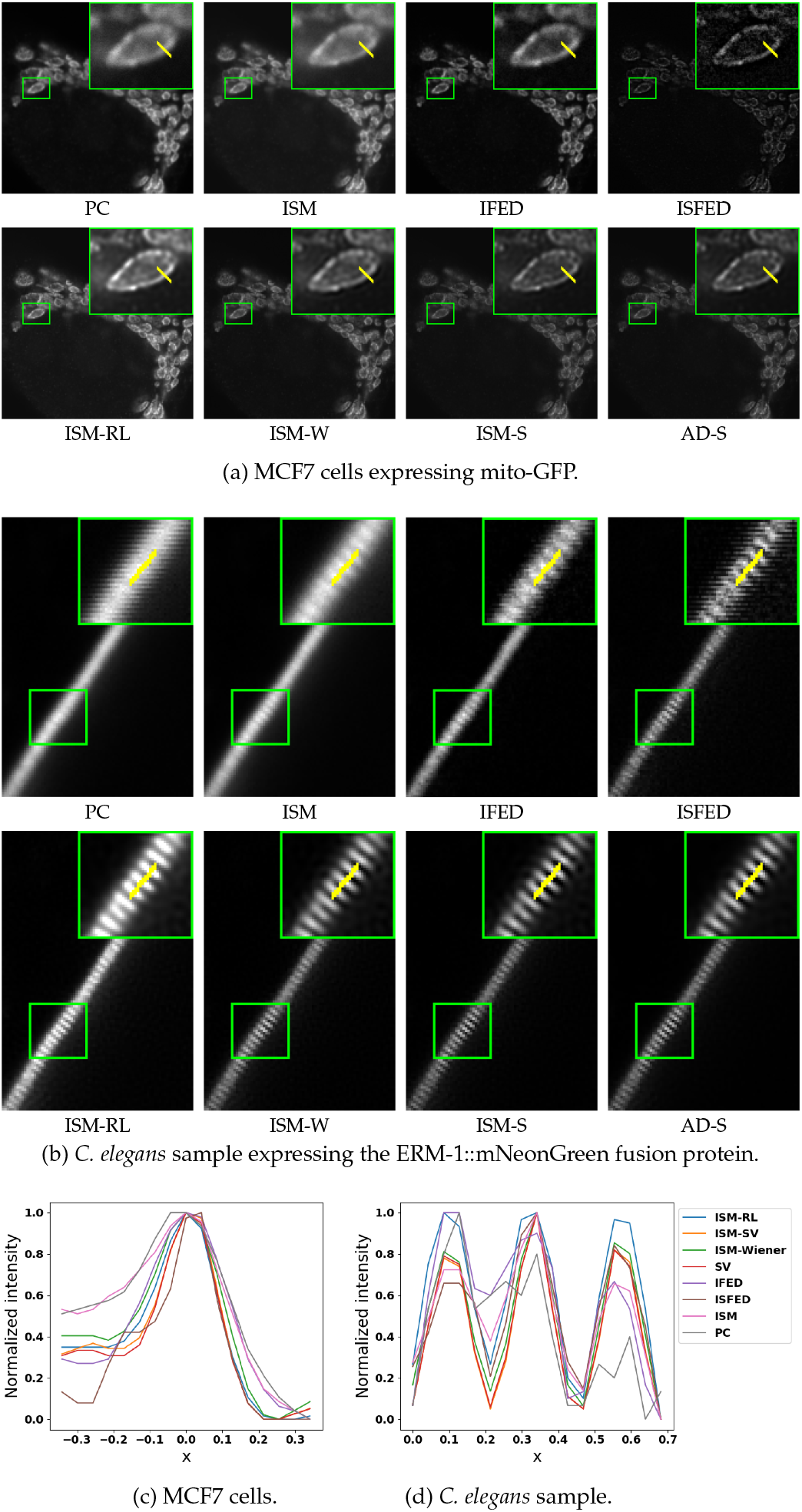
Reconstruction results obtained with the eight methods. (a) high resolution images depicting MCF7 cells expressing mito-GFP (labelling mitochondria). (b) High-resolution images depicting *C. elegans* sample expressing the ERM-1::mNeonGreen fusion protein. (c)-(d) Intensity profiles along the yellow line in (a) and (b). The *x* axes are graduated as the distances to the line centers in microns.

Finally, we performed another experiment using a *C. elegans* intestinal microvilli sample to quantitatively evaluate the methods sensitivity to SNR. The images have been captured with four different pixel dwell times, corresponding to four different acquisition speeds. For high pixel dwell time, the acquisition speed is low and the SNR is then high; when the pixel dwell time decreases, the acquisition speed increases and the SNR decreases. Figure 5 (left) shows the PC images for two acquisition speeds (speeds #1 and #4) in which the SNR decreases when the speed increases. We applied the eight reconstruction methods in each case and we evaluated the intensity contrast (in percentage) between microvilli (see plots in Fig. 5 (right). We notice that the contrasts obtained with the PC and ISM methods are below 26.5%, suggesting that we cannot distinguish the microvilli. With the IFED method, we can distinguish the microvilli only for speed #1. With the ISFED method, the microvilli can be separated for speeds #1 and #2. Unlike the linear methods, the deconvolution methods allows to resolve the microvilli for the four scanning speeds. As before, the AD-S method produced the best contrasted images when the SNR decreases.

**Fig. 5.**
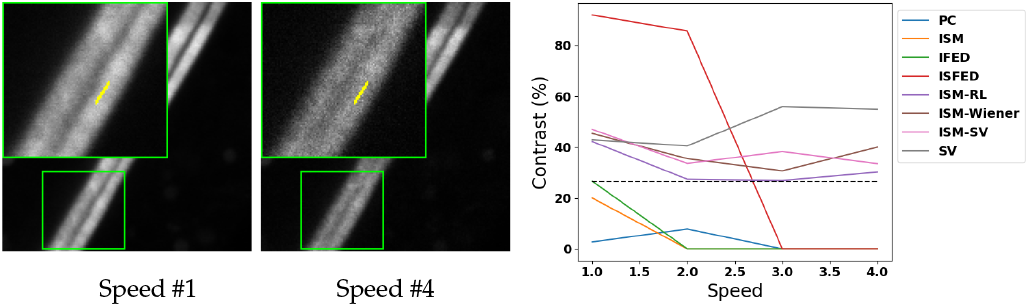
(left) Images of a *C. elegans* sample expressing the ERM-1::mNeonGreen fusion protein using two different scanning speeds from slow (Speed #1) to fast (Speed #4). (Right) Plots of the intensity contrasts in percentage between microvilli for each method.

In this Letter, we compared the main techniques to reconstruct a high resolution confocal image from array detectors, with a new array detector deconvolution method. Our AD-S method outperforms the state-of-the-art methods in terms of resolution and robustness to noise. The performance is still high provided the sub-detectors are satisfyingly co-registered and the PSF is approximately modeled by a Gaussian model. In this Letter, the comparison was performed on 2D images, but all the eight methods have been implemented to process 2D and 3D images. As for the computational performance, the linear methods take less than one second of computing time (core i7 CPU) to process a 512 × 512 image and 32 sub-detectors. The deconvolution methods are twice as slow (about 2 seconds).

## Funding

France-BioImaging ANR-10-INBS-04-07.

## Disclosures

The authors declare no conflict of interest.

## Data availability

Data acquisition was performed at the MRic Microscopy Rennes Imaging Center (France) with a confocal (Zeiss LSM 880) with airyscan microscope. The underlying results presented in this Letter may be obtained from the authors upon reasonable request.

## Software availability

The software (Fiji plugin) that gathers the tested methods is available here: https://team.inria.fr/serpico/airyscanj/.

## Supplemental document

See Supplement for supporting content.

## SUPPLEMENTARY NOTE

### SUPPLEMENTARY FIGURES

**Supplementary Fig. 1:**
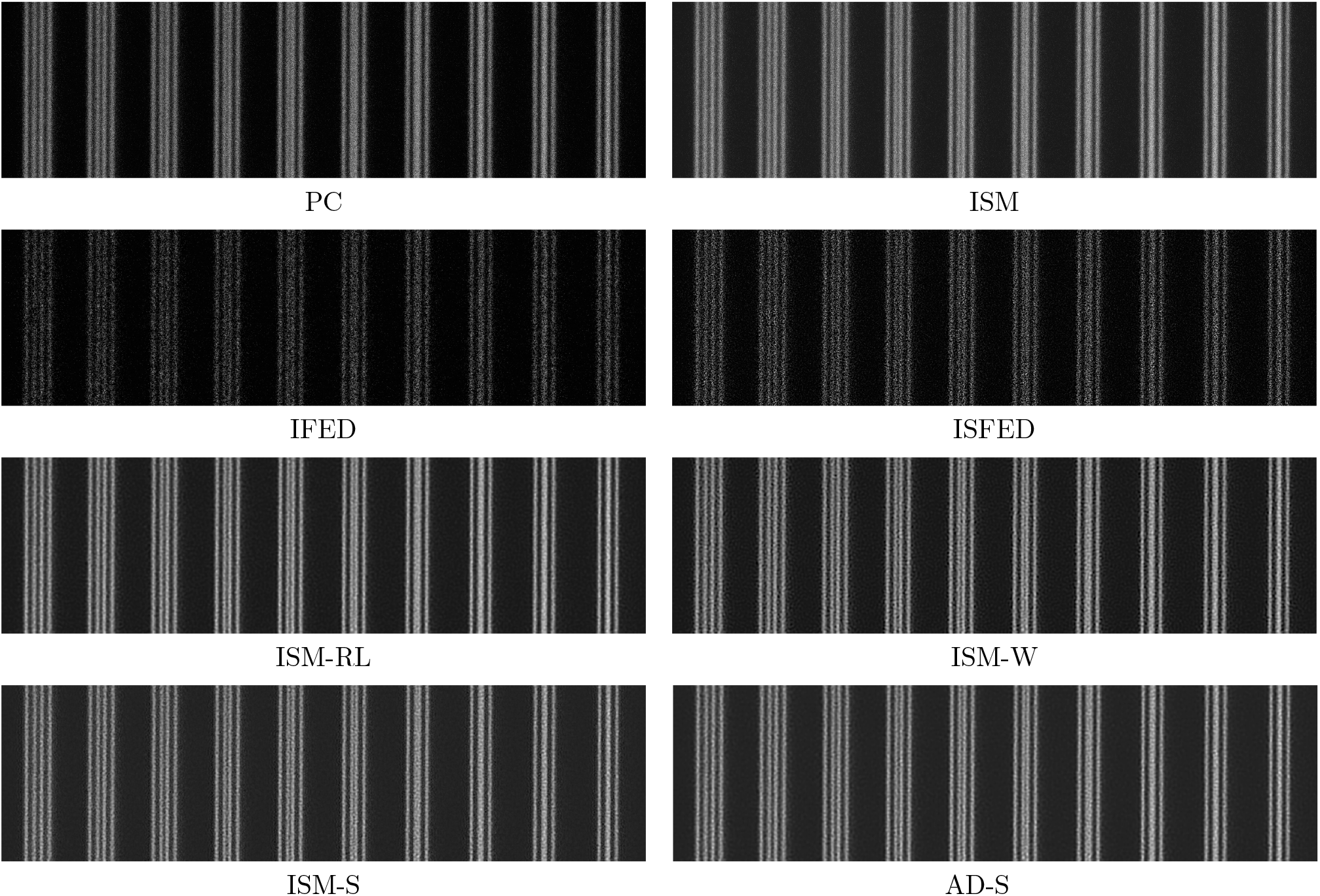
Reconstruction results obtained calibration slide containing vertical fluorescent lines spaced by 550, 500, 450, 400, 350, 300, 250, 200, 250, and 100 nm.

**Supplementary Fig. 2:**
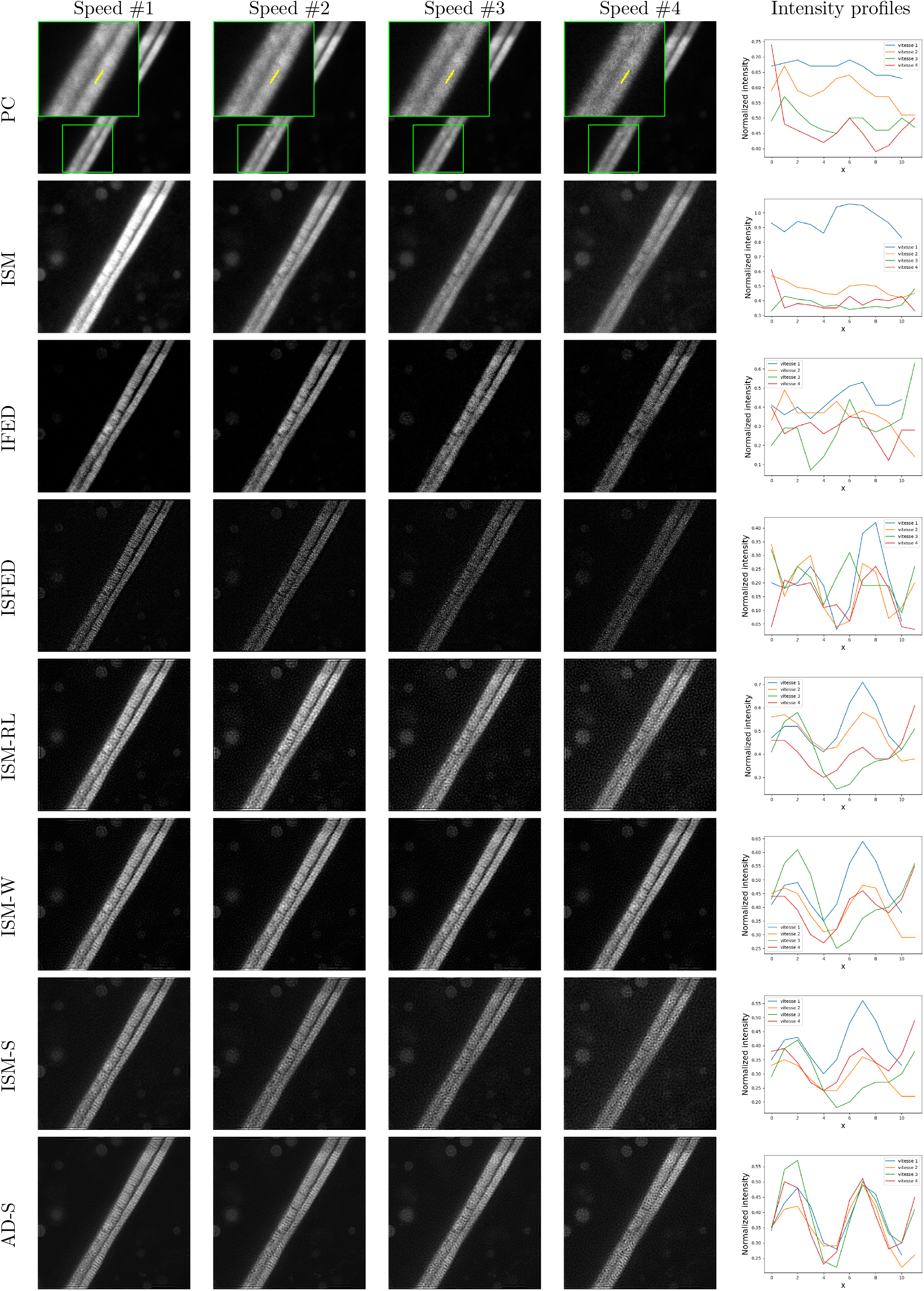
Images of a *C. elegans* sample expressing the ERM-1::mNeonGreen fusion protein using using two different scanning speeds from slow (Speed #1) to fast (Speed #4). The profile line location are marked with green patches on the PC images. The intensity profiles along the yellow lines are plotted in the right column. The x axes are graduated as the distances to the line centers in microns.

## APPENDIX

### A. Estimation of the optimal subtraction factor *∊*

The IFED and ISFED methods are based on the subtraction of two sets of detectors. To generalize the idea, we define the reconstructed image *û*(*x*) at pixel *x* ∈ Ω as:

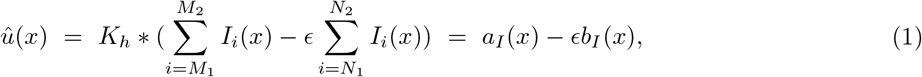

where *h* is the bandwidth of the convolution filter *K* and * is the convolution operator. In what follows, 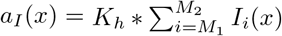 is the sum of the detectors with indexes between *M*_1_ and *M*_2_, and 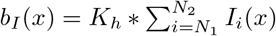 is the sum of the detectors with indexes between *N*_1_ and *N*_2_, and *∊* > 0 is a constant. In the case of *IFED*, *M*_1_ = 1, *M*_2_ = *N*, *N*_1_ = *N* + 1 and *N*_2_ = 32, and in the case of *ISFED*, *M*_1_ = 1, *M*_2_ = 32, *N*_1_ = 1 and *N*_2_ = 32. The bandwidth *h* is chosen very small to slightly reduce noise while preserving high frequencies.

Let us consider a neighborhood Ω(*x*) ⊂ Ω centered at point *x* and let us denote *n* = |Ω(*x*)| the number of pixels in Ω(*x*), assumed to be constant in the image image Ω. We propose to estimate the subtraction factor *∊* by using the local Stein’s unbiased risk estimate (*SURE*) of *û*(*x*) by considering *I*_1_ as the reference image:

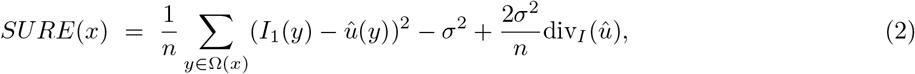

where *σ*^2^ is the variance of the assumed white Gaussian noise and

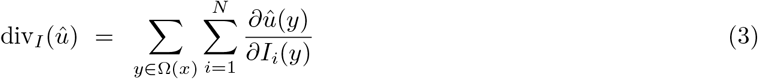

is the divergence of *û*. Since *a_I_*(*y*) and *b_I_*(*y*) have simple forms, it follows that:

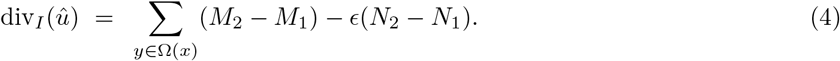

The *SURE* risk is then written as:

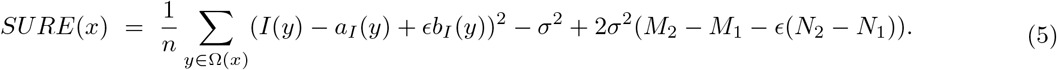

Furthermore, we define the empirical *SURE* as the sum of local *SURE* as:

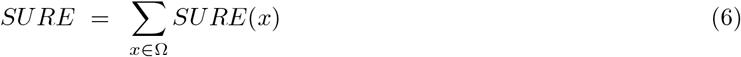

which is an unbiased estimator of the Mean Square Error (MSE), that is:

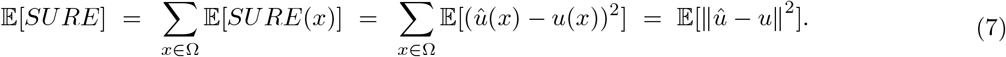

It is worth noting that, when the neighborhood Ω(*x*) is reduced to a single pixel *x*, the empirical SURE is nothing else that the conventional *SURE* on domain Ω. It follows that solving 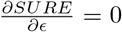 gives the following closed-form solution for *∊*:

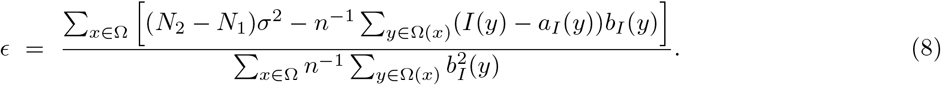

## Footnotes

1 In the case of *IFED*, *M*_1_ = 1, *M*_2_ = *N*, *N*_1_ = *N* + 1 and *N*_2_ = 32; in the case of *ISFED*, *M*_1_ = 1, *M*_2_ = 32, *N*_1_ = 1 and *N*_2_ = 32.

## Notes

https://team.inria.fr/serpico/airyscanj/

### Competing Interest Statement

The authors have declared no competing interest.

